# Attentional reorientation along the meridians of the visual field: are there different neural mechanisms at play?

**DOI:** 10.1101/816165

**Authors:** Simon R. Steinkamp, Simone Vossel, Gereon R. Fink, Ralph Weidner

## Abstract

Hemispatial neglect, after unilateral lesions to parietal brain areas, is characterized by an inability to respond to unexpected stimuli in contralesional space. As the visual field’s horizontal meridian is most severely affected, the brain networks controlling visuospatial processes might be tuned explicitly to this axis. We investigated such a potential directional tuning in the dorsal and ventral frontoparietal attention networks, with a particular focus on attentional reorientation. We used an orientation-discrimination task where a spatial pre-cue indicated the target position with 80% validity. Healthy participants (n = 29) performed this task in two runs and were required to (re-)orient attention either only along the horizontal or the vertical meridian, while fMRI and behavioral measures were recorded. By using a General Linear Model for behavioral and fMRI data, Dynamic Causal Modeling for effective connectivity, and other predictive approaches, we found strong statistical evidence for a reorientation effect for horizontal and vertical runs. However, neither neural nor behavioral measures differed between vertical and horizontal reorienting. Moreover, models from one run successfully predicted the cueing condition in the respective other run. Our results suggest that activations in the dorsal and ventral attention networks represent higher-order cognitive processes related to spatial attentional (re-)orientating that are independent of directional tuning.

## 1 Introduction

We are constantly exposed to an almost infinite amount of incoming sensory information. However, our brain’s capacities to process new data are limited. An efficient selection of important over unimportant information is, therefore, critical to ensure efficient information processing. This selection process, in which salient features of the sensory environment, as well as our internal goals and preferences, are considered, is commonly referred to as selective attention.

The allocation of attentional resources is controlled by neural structures that are thought to be organized in two distinct but interacting frontoparietal networks (Corbetta, Patel, & Shulman, 2008; Corbetta & Shulman, 2011; Vossel, Geng, & Fink, 2014).

The top-down guided (i.e., voluntary) orienting of attention involves a bilaterally organized dorsal frontoparietal network, encompassing the intraparietal sulcus (IPS) and the frontal eye-fields (FEF). Converging evidence from functional imaging and transcranial magnetic stimulation (TMS) suggests that these regions may modulate the activity in sensory (e.g., visual) cortices to prioritize the processing of stimuli at specific locations in space (e.g., Bressler, Tang, Sylvester, Shulman, & Corbetta, 2008; Hung, Driver, & Walsh, 2011; Ruff et al., 2008; Vossel, Weidner, Driver, Friston, & Fink, 2012).

Unexpected or very salient stimuli may interrupt our current top-down guided focus of attention (Simons, 2000), initiating a redistribution of processing resources. In this case, the allocation of attention is guided in a bottom-up fashion, meaning that it is primarily based on external stimulus features. The ventral frontoparietal attention network supposedly regulates this bottom-up control of attention. A central node within this network is the temporoparietal junction (TPJ), which has been suggested to be the driving force for attentional reorienting (Corbetta et al., 2008). The ventral network further consists of the inferior and the middle frontal gyrus (IFG, MFG) and is typically described as being strongly lateralized to the right hemisphere (Corbetta & Shulman, 2011). Recent studies, however, show that left TPJ is also involved in controlling spatial attention (Beume et al., 2017; Silvetti et al., 2016).

Unilateral lesions following a stroke can lead to an inability to allocate attention to the visual field contralateral to the lesion (Halligan, Fink, Marshall, & Vallar, 2003) - a phenomenon often referred to as hemispatial neglect. Neglect is more frequent and severe following right-hemispheric lesions and causes symptoms predominantly in contralesional space (Karnath, Rennig, Johannsen, & Rorden, 2011). This lateralization suggests a unique role for orienting and reorienting attention along the horizontal meridian and hence motivated research with a focus on that particular spatial dimension. Attentional orienting along the vertical meridian on the other hand seems understudied, despite the fact that there is also evidence for a vertical component in hemispatial neglect (Cappelletti, Freeman, & Cipolotti, 2007) and that cases of vertical neglect of the upper visual field after bilateral lesions to the inferior temporal lobes have been reported (Shelton, Bowers, & Heilman, 1990). Vertical neglect commonly affects the lower left visual field after right hemispheric lesions (Cazzoli, Nyffeler, Hess, & Müri, 2011; Müri, Cazzoli, Nyffeler, & Pflugshaupt, 2009; Pitzalis, Spinelli, & Zoccolotti, 1997). The extent of horizontal and vertical neglect along the meridians seems to be additive, becoming more pronounced at oblique positions (i.e., lower left visual field), which has also been observed for pseudo-neglect in healthy participants (Nicholls, Mattingley, Berberovic, Smith, & Bradshaw, 2004). Thus, the allocation of attention along the two meridians may rely on distinct neural mechanisms.

However, it remains unclear if the brain regions controlling shifts of spatial attention are tuned to specific spatial directions or if they constitute a uniform system with no particular spatial preference (i.e., directional tuning). Several attempts have already been made to disentangle the neural mechanisms underlying vertical as compared to horizontal attentional orienting. The evidence coming from different neuroimaging studies, however, is inconclusive about the brain regions involved. On the one hand, orienting attention along a horizontal relative to a vertical axis activated the lingual and right precentral gyrus, whereas orienting attention in a vertical dimension involved more pronounced activation in the precuneus, medial frontal cortex, anterior cingulate, and cerebellum (Mao, Zhou, Zhou, & Han, 2007). Furthermore, ventral medial prefrontal cortex, cuneus, and lingual gyrus have been reported to be more involved in horizontal as compared to vertical antisaccades (Lemos et al., 2017), and left FEF and left superior temporal gyrus are more related to vertical relative to horizontal prosaccades (Lemos et al., 2016). Several other studies could not find any evidence for differences between horizontal and vertical attentional processes (Fink, Marshall, Weiss, & Zilles, 2001; Macaluso & Patria, 2007).

Therefore, the goal of the present fMRI study was to clarify the involvement of attentional control areas in reorienting attention along the vertical and horizontal meridian. To this end, both blood oxygenation level dependent (BOLD) amplitudes and measures of effective connectivity were employed. We used a variant of Posner’s spatial cueing paradigm (Posner, 1980) in which participants had to indicate the orientation of a Gabor patch via button presses while ignoring distractor stimuli at other locations. A pre-cue (arrow) indicated the most likely target location. Spatial reorienting of attention was induced by presenting invalid cues in 20% of the trials. The experiment involved two runs that differed about the spatial direction of attentional orienting and reorienting. In these two different runs, cues and targets were presented either along the vertical or the horizontal meridian of the visual field. Potential differences in attentional processing along the vertical or horizontal meridian concerning the BOLD-amplitudes were expected to induce a main effect of direction (horizontal, vertical), or an interaction between direction and cueing (valid, invalid cueing) in the standard general linear model (GLM) analysis of the fMRI data. Furthermore, vertical and horizontal reorienting of attention were expected to generate differential effective connectivity patterns in the activated brain areas in an analysis using Dynamic Causal Modelling (DCM, Friston, Harrison, & Penny, 2003).

## 2 Methods

### 2.1 Participants

We recruited 29 right-handed participants (Edinburgh handedness inventory (Oldfield, 1971), M = 0.86, SD = 0.14) with corrected to normal vision, who gave written informed consent. One participant had to be excluded from both behavioral and fMRI analysis due to noncompliance with the task. Another participant was excluded only from further fMRI analysis due to excessive head movements (translation > 3 mm); however, the participant’s behavioral data were included in further analysis. The remaining 28 participants (15 female) were between 21 to 39 years (M = 25, SD = 3) old. The ethics board of the German Psychological Association had approved the study. Participants were compensated with 15€ per hour.

### 2.2 Experiment

Participants performed a spatial cueing paradigm inside a 3 Tesla TRIO MRI scanner (Siemens, Erlangen). Stimuli were displayed on a screen that was mounted at the end of the scanner’s bore and could be seen by the participant via a mirror (245 cm distance). The mirror was mounted on top of a 32 channel head coil. Participants’ task throughout the experiment was to report the orientation (horizontal 90° or vertical 0° rotation) of a target stimulus (Gabor patch, diameter 1° visual angle) using button presses of their left and right index fingers. Participants were instructed to continually fixate a diamond in the screen’s center (0.5° wide). Next to the central diamond, empty boxes (1° wide) were presented in all four cardinalities throughout the experiment with their centers at 4° eccentricity. Each trial began with an alerting signal, a 500 ms brightening of the diamond’s center, followed by a spatial cue (duration: 200 ms) after 1000 ms. Brightening and widening of one of the central diamond’s corners served as a symbolic cue (arrowhead), indicating the most likely upcoming target location with 80% probability. We informed the participants about the cue validity during the task instructions. After a variable interval of 400 or 600 ms, the target stimulus appeared (duration: 250 ms) in the cued box (valid trial) or in the box opposite to the cue (invalid trial). Distractor stimuli were presented in the remaining three boxes for the same duration as the stimulus. Distractors were created by superimposing two Gabor patches, which were rotated by 45° and 135°. The resulting pattern matched the target stimulus in intensity and contrast (see Figure 1). The inter-trial interval separating subsequent trials was either 2.0 s, 2.7 s, 3.2 s, 3.9 s, or 4.5 s with equal probability. Trials were presented in two subsequent runs, with a short break in between. In one run, cues pointed only to left or right, and target stimuli were only presented along the horizontal meridian. In the other run, cues pointed only upwards or downwards, and the target only appeared in the upper or lower box (i.e., on the vertical meridian). Before each run, participants completed 20 practice trials with immediate feedback regarding accuracy. Each run consisted of 5 blocks, each comprising 32 valid and 8 invalid trials. The 8 possible target properties (position left/right or up/down, left/right response finger, 400/600 ms SOA) were presented with equal probability in each block. Trial order in each block, however, was fully randomized. The order of horizontal and vertical runs and the response mapping (left or right finger for horizontally oriented stimuli) were counterbalanced across participants. Between the different blocks, a 10 to 13 s break period was included. Before the actual spatial cueing paradigm, participants also completed a separate short training to get used to the response mapping between stimulus orientation and response fingers. Here, sixty target stimuli appeared rapidly in the screen’s center, and participants had 500 ms time to respond. Immediate feedback was given, and the percentage of correct responses was continuously presented. Recording of responses and stimulus presentation were controlled with PsychoPy (version 1.85.3, Peirce, 2007, 2008; Peirce et al., 2019).

**Figure 1.**
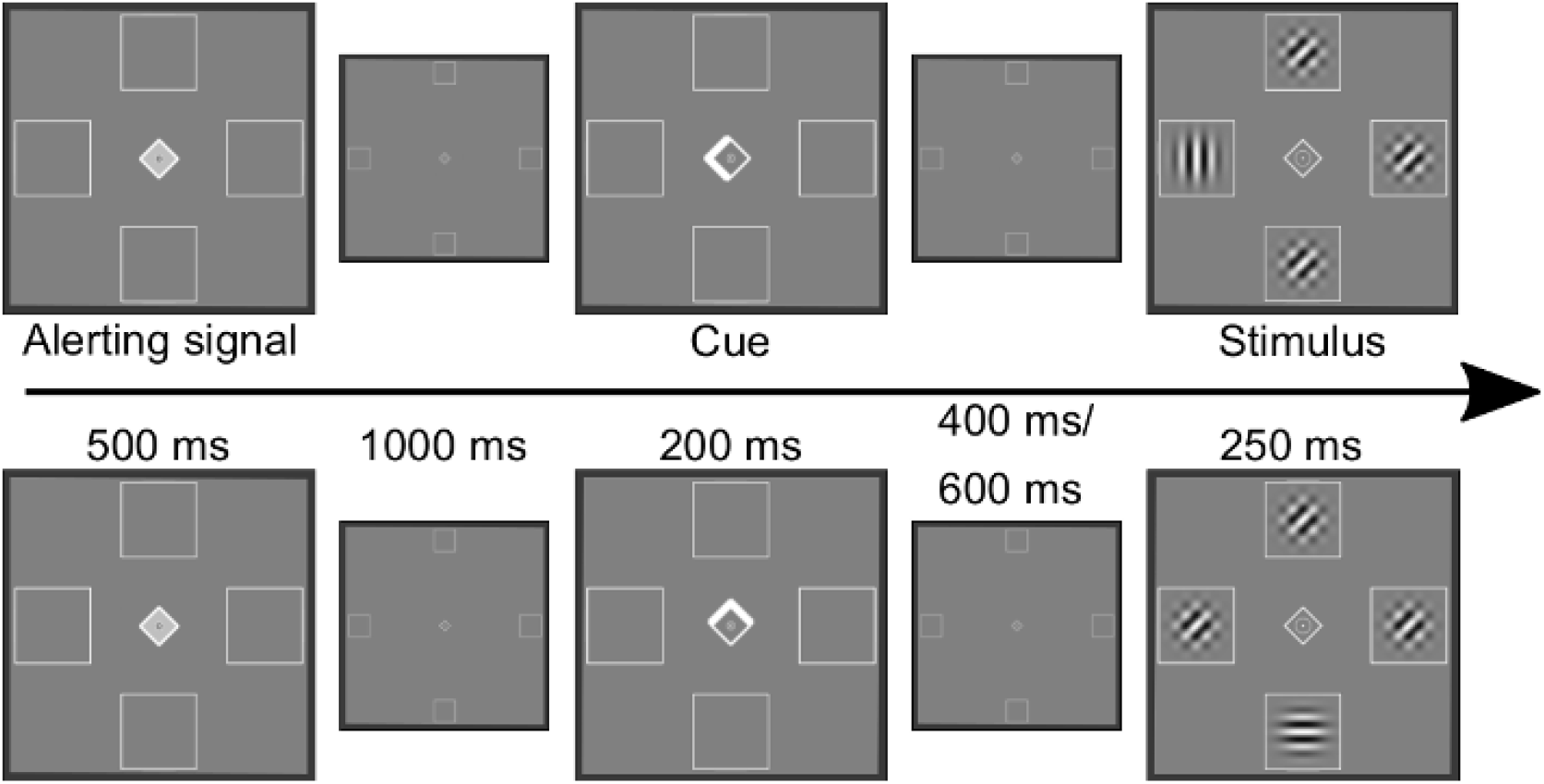
One trial for each run of the cued attention task. In the upper row, a valid trial in the horizontal session is displayed, in the lower row, an invalid trial of the vertical session. Displays for the alerting signal, cue, and stimulus presentation were enlarged for better presentation. The smaller displays show the stimulus presentation in the correct right proportion. Participants were told always to fixate the center of the screen. Their task was to press a button corresponding to the orientation of the target stimulus (vertical or horizontal Gabor patches).

### 2.3 Behavioral analyses

We used a two-step procedure to test for differences in reaction times and error rates between the vertical and horizontal runs and the effects of valid and invalid cueing. The resulting 2 (cueing: valid/invalid cues) × 2 (direction: horizontal/vertical) design was subjected to a Bayesian implementation of an analysis of variance (BF_ANOVA), treating participants as random factors (Rouder, Morey, Speckman, & Province, 2012). The Bayes factors (BF) for different models representing the possible combinations of factors were calculated using the BayesFactor package (version 0.9.12-4.2, Morey & Rouder, 2018) implemented in R (version 3.5.1, R Core Team, 2018), using default settings (‘medium’ scaling factor on the JSZ-prior and 10000 iterations of the MCMC algorithm). The BF_10_ in favor of the model (H_1_) was calculated by dividing the model’s posterior probability by the posterior probability of a null model (grand mean plus random factors, H_0_). Additionally, we compared the model with the highest BF_10_ against all the other models (main effects and interaction). Following standard conventions, a BF_10_ > 3 is regarded as positive evidence and a BF_10_ > 10 as strong evidence in favor of H_1._ A BF_10_ < 0.33 is then seen as positive evidence and a BF_10_ < 0.1 as strong evidence in favor of the null hypothesis (Jarosz & Wiley, 2014).

The error rates were calculated for each participant by taking the mean of incorrect and missed responses for each direction (horizontal/vertical) and cueing condition (valid/invalid). Reaction times were defined as the median response times for each direction and cueing condition. Before the calculation of the median, we removed the error and post-error trials (to account for post-error slowing), missed responses, trials with response times faster than 200 ms, and response times exceeding the 75% quartile + 1.5 * interquartile range criterion.

### 2.4 FMRI

We obtained 557 T2* weighted images per run using an echo planar imaging (EPI) sequence (time of repetition (TR) 2.2 s; echo time (TE) 30 ms; flip angle 90°). Each image consisted of 36 transverse slices (recorded in an interleaved and ascending manner), with a voxel size of 3.1 × 3.1 × 3.3 mm and 3 mm slice thickness (field of view 200 mm). We manually discarded the first 5 images of each run to account for T1 equilibrium artifacts. In addition to the BOLD images, we obtained a structural T1 anatomical image for each participant.

The following description of the preprocessing was automatically generated (see http://fmriprep.readthedocs.io/en/1.1.1/workflows.html) and minimally adapted.

The preprocessing of functional and anatomical data was performed using FMRIPREP version 1.1.1 (Esteban et al., 2018, 2019, RRID:SCR_016216), a Nipype (RRID:SCR_002502, Gorgolewski et al., 2011, 2017) based tool, run as a docker-image. Each T1-weighted volume (T1w) was corrected for intensity non-uniformity using N4BiasFieldCorrection v2.1.0 (Tustison et al., 2010) and skull-stripped using antsBrainExtraction.sh v2.1.0 (using the OASIS template). Spatial normalization to the ICBM 152 Nonlinear Asymmetrical template version 2009c (Fonov, Evans, McKinstry, Almli, & Collins, 2009, RRID:SCR_008796) was performed through nonlinear registration with the antsRegistration tool of ANTs v2.1.0 (Avants, Epstein, Grossman, & Gee, 2008, RRID:SCR_004757), using brain-extracted versions of both T1w volume and template. Brain tissue segmentation of cerebrospinal fluid (CSF), white-matter (WM), and gray-matter (GM), was performed on the brain-extracted T1w using fast (Zhang, Brady, & Smith, 2001, FSL v5.0.9, RRID:SCR_002823).

Functional data were slice time corrected using 3dTshift from AFNI v16.2.07 (Cox, 1996, RRID:SCR_005927) and motion-corrected using mcflirt (FSL v5.0.9, Jenkinson, Bannister, Brady, & Smith, 2002). “Fieldmap-less” distortion correction was performed by co-registering the functional image to the same-subject T1w image with intensity inverted (Wang et al., 2017) constrained with an average fieldmap template (Treiber et al., 2016), implemented with antsRegistration (ANTs). This procedure was followed by co-registration to the corresponding T1w using boundary-based registration (Greve & Fischl, 2009) with 9 degrees of freedom, using flirt (FSL). Motion correcting transformations, field distortion correcting warp, BOLD-to-T1w transformation, and T1w-to-template (MNI) warp were concatenated and applied in a single step using antsApplyTransforms (ANTs v2.1.0) using Lanczos interpolation.

Frame-wise displacement (Power et al., 2014) was calculated for each functional run using the implementation of Nipype.

Many internal operations of FMRIPREP use Nilearn (Abraham et al., 2014, RRID:SCR_001362), principally within the BOLD-processing workflow. For more details of the pipeline, see http://fmriprep.readthedocs.io/en/1.1.1/workflows.html.

Additional spatial smoothing of the functional images was performed in SPM12 (version 7219, Friston, 2007) implemented in MATLAB 2016b (The MathWorks, Inc., Natick, Massachusetts, United States), using an 8 mm FWHM Gaussian kernel.

### 2.5 Analyses of imaging data

The first level statistical analysis of the data was performed using SPM12. For group-level analysis, we used the statistical non-parametric mapping (SnPM) toolbox (version 13.1.07, Nichols & Holmes, 2002). At the single-subject level, we modeled both runs in the same design matrix using an event-related design (i.e., a stimulus duration of 0), with run-specific intercepts and confounds. As regressors of interest, we used the target onsets of the two cueing-conditions and the four possible target positions. This resulted in eight different regressors for invalid left (iL), invalid right (iR), valid left (vL), valid right (vR), as well as invalid down (iD), invalid up (iU), valid down (vD), and valid up (vU) trials. For each run, up to two additional regressors were added. One regressor was used to account for error and post-error trials and another to account for outlier trials (please, see the behavioral analysis for the definition of outliers). The regressors’ onsets were convolved with a canonical hemodynamic response function (HRF). The six movement parameters calculated during realignment and the frame-wise displacement were included in the model as confounds. A cosine set accounting for drifts and high-pass filtering was applied following the SPM12 defaults.

We investigated five planned contrasts: (1) The main effect of all invalid versus valid trials ((iL + iR + iD + iU) – (vL + vR + vD + vU)), (2 & 3) two contrasts for direction-specific cueing effects: horizontal reorientation (iL + iR) – (vL + vR) and vertical reorientation (iD + iU) – (vD + vU), (4) a contrast for the main effect of direction ((iL + iR + vL + vR) – (iD + iU + vD + vU)), and (5) a contrast for the interaction of cueing and direction ((iL + iR – vL – vR) – (iD + iU – vD – vU)). Additional four tests were performed to show the effects of attentional and perceptual modulation in the visual areas by valid targets. These tests were performed separately for each visual field (vL > vR; vR > vL; vD > vU; vU > vD).

Group level t-maps were then calculated for each contrast using one-sample permutation t-tests (25000 permutations, no variance smoothing) with a predefined cluster forming threshold of p < 0.001 uncorrected (SnPM: fast option). We report the results of thresholded t-maps, using a significance cut-off of p < 0.05 (FWE corrected at the predefined cluster level). An overview of global and local maxima was created using the function “get_clusters_table” implemented in the Python package Nistats (version 0.0.1b, Abraham et al., 2014).

### 2.6 VOI Analyses

As we did not find any significant differences in BOLD amplitudes between horizontal and vertical directions, and no significant activations for the interaction of direction and cueing (valid/invalid), we conducted a more sensitive post-hoc VOI based analyses. Here, we probed bilateral TPJ, FEF, and IPS, which are key regions of the ventral and dorsal attention networks. The global or local maxima corresponding to the six regions of the condition main effect (Table 1) defined the seed coordinates. These were passed to Nilearn’s (version 0.4.2) “NiftiSpheresMasker” function (without standardization and detrending)—implemented in Python 3.7—to extract the mean beta values of the eight regressors of interest using an 8 mm sphere, masked by the thresholded t-map of the main cueing effect (1).

**Table 1:**
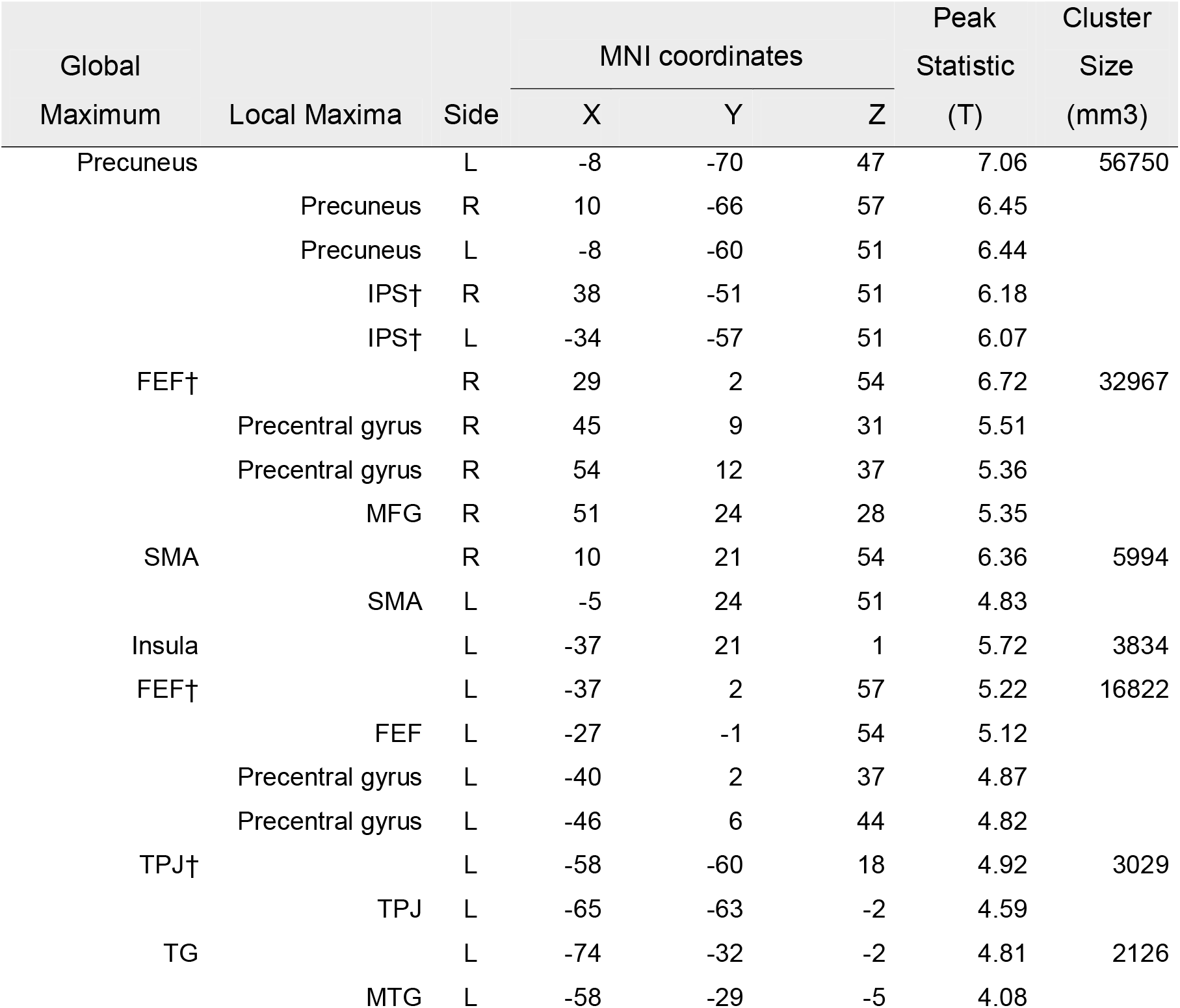
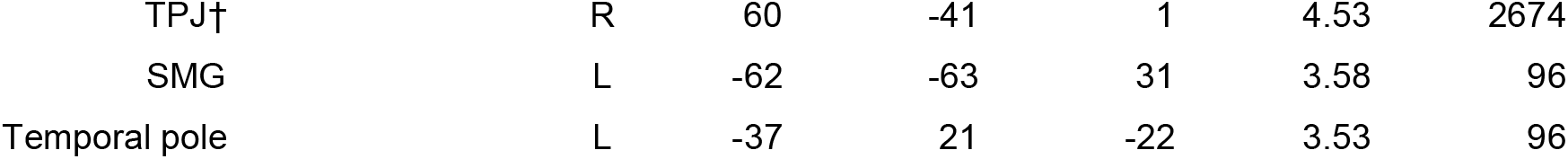
Cluster coordinates for reorienting across horizontal and vertical runs. Global and up to four local maxima’s coordinates and peak t-statistics of the thresholded statistical maps. Coordinates annotated with a dagger (†) were included in the further VOI-based and DCM analyses. Rounded MNI coordinates and cluster sizes were estimated using Nistats’ get_cluster_table. *IPS – intraparietal sulcus, FEF – frontal eye fields, MFG – middle frontal gyrus, SMA – supplementary motor area, TPJ – temporoparietal junction, TG – temporal gyrus, MTG – middle temporal gyrus, SMG – supramarginal gyrus*.

The mean beta values were averaged over visual fields to obtain values for the horizontal and vertical directions. For example, the beta value used for left IPS during invalid horizontal trials consisted of the average extracted beta values from iL and iR. For each VOI, we analyzed whether direction or interaction effects were present using BF_ANOVAs, with the participant as the random factor. We followed the rationale described for the analysis of the behavioral data.

In addition to the BF_ANOVAs, we used logistic regression to test whether brain activity differences between cueing-conditions of one direction were predictive for the cueing effect in the respective other direction. We again used the mean betas of each participant in the six VOIs for each of the 8 regressors (iL, iR, iD, iU, vL, vR, vD, vU), this time not collapsing along meridians. Then, we tested whether BOLD amplitude patterns in the six VOIs of the horizontal run (iL, iR, vL, vR) were similar enough to differentiate valid and invalid trials of the vertical run (iD, iU, vD, vU), and vice versa. This was done using logistic regression implemented in scikit-learn (version 0.20.0, Pedregosa et al., 2011). The logistic regression’s performance was first estimated on a per run basis using nested cross-validation. Each run’s data was split into fives, so that every split served as test-data once. For each round, the remaining splits served as training data and were again subjected to 5-fold cross-validation to find the best regularization parameter C in the range [10^−4^, 10^−3^ …, 10^3^, 10^4^]. The regularization parameter that achieved the highest average accuracy in the inner cross-validation loop was used to refit the logistic regression on all of the training data. The run-based model performance was then defined as the average accuracy over the splits. A similar approach was used to estimate generalized performance, where 5-fold cross-validation was used on one run to find the best parameter C, and the accuracy was calculated for the predictions made on the other run.

As a performance measure, we used permutation tests by shuffling the class-labels (valid or invalid trials), refitting the logistic regression and then recalculating the accuracies (1000 permutations). The permutation P-value then represents the proportion of accuracy scores that were higher in the random condition than in the original (Ojala & Garriga, 2010).

### 2.7 DCM Analyses

In addition to differences in BOLD amplitudes, we were interested in the cueing-dependent effective connectivity patterns in the horizontal and vertical runs. To estimate effective connectivity, we used bilinear DCM (DCM 12, revision 6755, in MATLAB 2016b). DCM is a state-space model described by a directed graph to infer the cortical dynamics in time between brain regions. Each node in the graph is defined by a brain region and represents its neural activity. The edges represent how nodes influence each other. This approach leads to a generative model that, once inverted, can be used to simulate neural activity in the network. Therefore, it can be used to investigate how network dynamics would differ in the presence of different inputs. The state-change equation of neural states in DCM is described by equation 1 (Friston et al., 2003).

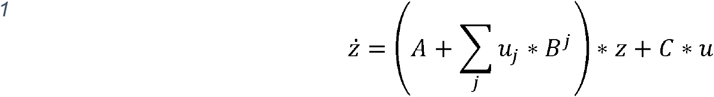

The change in the hidden neural states ż is described by the fixed connectivity matrix *A*, which represents the coupling between brain regions in the absence of exogenous modulations (*u*). The coupling can be modulated by the *j* exogenous inputs (*u*), which are represented by the parameters in the matrix *B* (the connections in *B* are a subset of *A*). Lastly, the driving input regions, which represent the direct changes of hidden states, is defined by the matrix *C*. As we were interested in how connection strength differs between invalid trials in the horizontal (*u*_*1*_) as compared to the vertical run (*u*_*3*_), we restricted our analysis to the parameters in the matrices *B*^*1*^ and *B*^*3*^. Matrices *B*^*2*^ and *B*^*4*^ were left empty, which means that connections were not modulated by valid trials (neither in the horizontal nor the vertical run). Since we were interested in investigating potential differences in reorienting of attention (invalid trials), we assumed that connectivity between brain regions in valid trials was the same for both runs (i.e., that all dynamics of valid trials were captured in the baseline connectivity described by the matrix *A*).

As DCM for fMRI describes the BOLD dynamics between brain regions, we extracted the time series from the same regions as in the VOI analysis. By modeling activity in the IPS and FEF, we captured the neural dynamics during valid trials in the dorsal attention network. Additionally, bilateral TPJ enabled us to model the potential influence of the ventral attention network onto dorsal regions (“circuit breaker”, Corbetta et al., 2008). Thereby, we limited the network analysis to the most representative regions of the classic models of visual spatial attention. Due to our GLM findings that brain activity during valid trials was highly similar in both vertical and horizontal runs and only modulated by invalid trials, we concatenated the time series (spm_concat) of both runs. For the estimation of our DCMs, we defined a new design matrix in SPM12 for each participant. The target onsets of invalid horizontal (iH), valid horizontal (vH), invalid vertical (iV), and valid vertical (vV) trials served as driving inputs to the DCM. As in the GLM analysis above, we included the seven motion parameters as nuisance regressors and added run specific intercepts (to center the time series of each run). To extract time series at participant-specific peaks, we calculated first level t-tests of activations in valid or invalid trials against baseline (vH + vV and iH + iV, thresholded at p < 0.05 uncorrected). The VOI coordinates highlighted in Table 1 were used as the starting points for the VOI extraction. The different locations served as the center of 12 mm spheres in which the participant’s nearest local maximum was selected. The new coordinates were then used as the center of an 8 mm sphere from which the first principle component of the BOLD signal was extracted. The spheres included only task activated voxels (threshold p < 0.05), and the time series were adjusted for the nuisance regressors and mean activity. We used the contrast of valid trials against baseline to select the VOIs for bilateral FEF and IPS, and the contrast invalid trials against baseline for the TPJ VOIs.

The underlying network structure describing the intrinsic coupling during the task (*A*) was defined by fully connected intra-hemispheric regions and inter-hemispheric connections of homologous regions. All nodes received all driving-inputs because visual input was carefully matched across conditions and visual areas were comparably activated. This also reflects graphical descriptions of the ventral and dorsal attention networks, where all of the six regions receive information from the visual areas (Vossel et al., 2012).

Hierarchical family-wise Bayesian model selection (BMS) implemented in the MATLAB VBA-toolbox (version: master/7ac4470b987796cf4ec9bfb275ab049d5aa97931, Daunizeau, Adam, & Rigoux, 2014) and subsequent Bayesian Model Averaging (BMA, implemented in SPM12) were used to find the best connections and parameters in *B*^*1*^ and *B*^*3*^ that best describe our data. In the first step, three model families were used to investigate whether modulations by invalid trials occurred only in the left, right, or in both hemispheres. This was done in order to investigate possible right lateralization of the ventral attention network. The second class of families was used to decide upon the direction of inter-hemispheric modulations between left and right IPS. The remaining modulations, which can be seen in Figure 2, then describe whether TPJ affects the dorsal attention network or vice versa. The model space was restricted so that at least one modulation between the dorsal and ventral attention network had to be present and that there were no bidirectional modulations. In total, we inverted 72 models per participant. The modulations by invalid trials in horizontal and vertical runs (i.e., in *B*^*1*^ and *B*^*3*^) were the same. Hence, while the connectivity parameters could differ, the overall modulation structure by invalid trials stayed the same. Finally, we used BMA on the winning model-family on a participant level, to get more reliable point estimates for the different connections.

**Figure 2.**
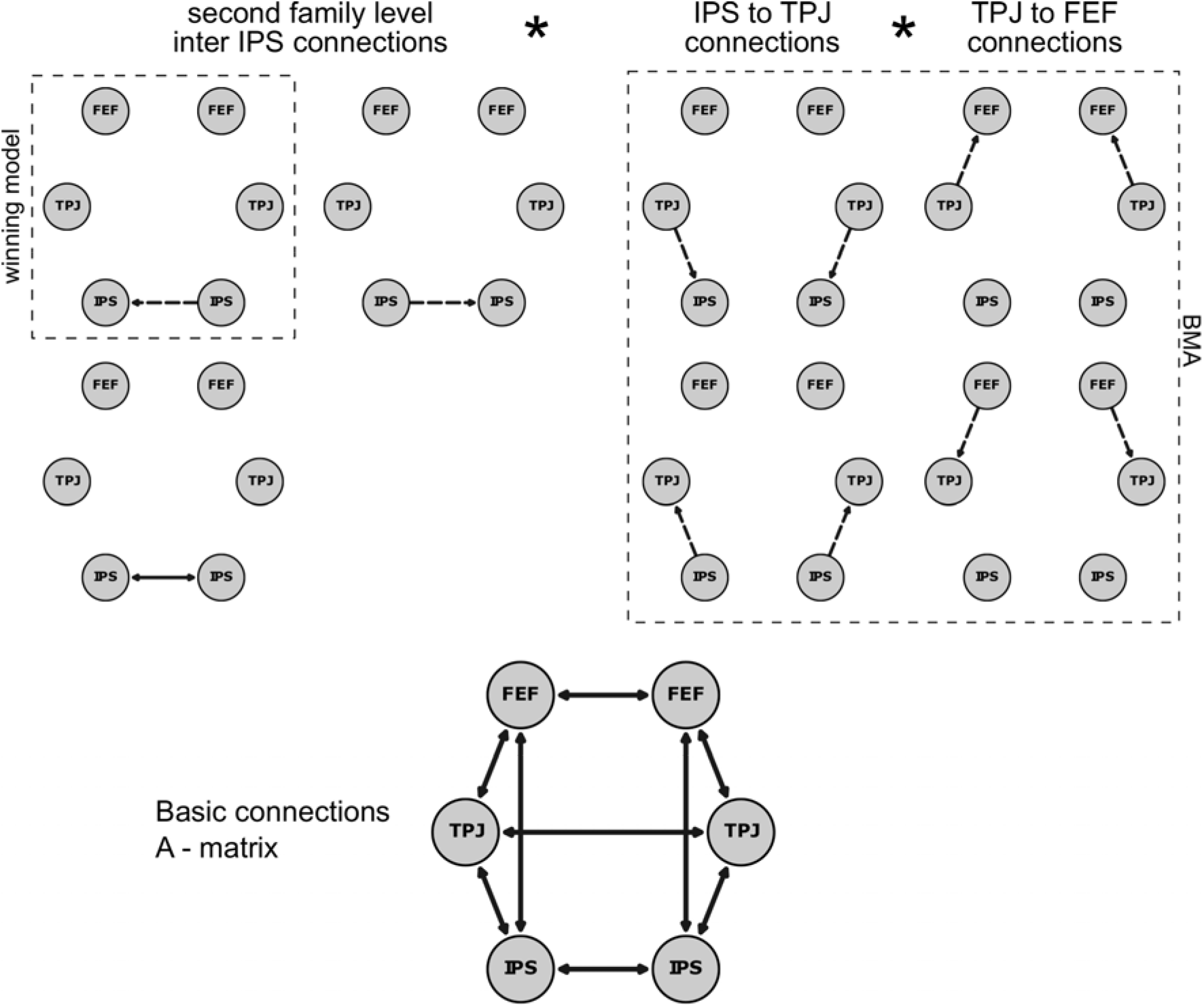
A schematic of the model space used in the fMRI analysis. The first model family comparison revealed that modulations between IPS and TPJ and FEF and TPJ were present in both hemispheres (models not shown). Hence, only the models of the bilateral family are shown. Dotted lines are used for unilateral and solid lines for bilateral connections. The second model family comparison favored models with a unidirectional connection from right IPS to left IPS (see left upper panel). The remaining model combinations based on connections between IPS-TPJ and FEF-TPJ were summarized using BMA. The model basis is shown in the lower part of the figure, indicating the fixed connections. *IPS – intraparietal sulcus; FEF – frontal eye-fields; TPJ – temporoparietal junction; BMA – Bayesian model average*.

The DCMs were created using custom MATLAB scripts, using mostly default settings for bilinear DCM. However, we used 36 instead of the 22 time steps in the discretization of the inversion function to account for slice time correction. Confounds, which were included in the DCM estimation, were manually added, so that temporal drifts, represented by a discrete cosine set, and confounds calculated during the participant’s SPM design matrix were included for each run separately.

We tested whether the modulation by invalid trials differed between the vertical and horizontal session by calculating the BF_10_ in favor of any difference between runs using Bayesian paired t-tests for each parameter pair in *B*^*1*^ and *B*^*3*^. Testing for differences in effective connectivity strength between runs, however, does not provide us with the full picture. For example, it remains unknown how the parameters interact as a whole within the network. Therefore, using the generative properties of DCM (and the BMA parameter estimates), we simulated the BOLD signal by swapping the inputs (*u*) between the horizontal and the vertical runs (i.e., iH (u_1_) ↔ iV (u_3_), vH (u_2_) ↔ vV (u_4_)). This approach allowed us to evaluate the specificity/generality of the parameters for horizontal and vertical reorienting of attention. If the model performance with the parameters of the respective other run is comparable to the original data, we can conclude that, regardless of specific parameter values, the neural processes of invalid trials are similar across runs.

The performance of the swapped model was compared against random models in which the onset timings of the impulses in *u* were kept, but the input streams (u_1_, u_2_, u_3_, u_4_) were assigned randomly. We report the proportion of participants with permutation P-values lower than p < 0.05 in the original and swapped conditions. The permutation P-values were calculated as the proportion of models where the root mean squared error (RMSE, eq.2) was larger in the original or swapped data than in 1000 sets of randomly generated data.

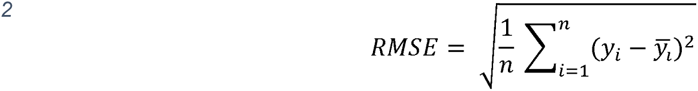

## 3 Results

### 3.1 Behavioral data

Participants’ reaction times in invalid trials were higher than reaction times in valid trials, both in the horizontal (invalid M = 718.32 ms, SD = 113.58 ms; valid: M = 674.50 ms, SD = 98.48 ms) and in the vertical run (invalid M = 723.68 ms, SD = 138.74 ms; valid: M = 667.04 ms, SD = 116.52 ms). A similar pattern was observed for error rates (Figure 3). In the horizontal run, error rates were higher for invalid compared to valid trials (invalid M = 4.46%, SD = 5.37; valid: M = 3.44%, SD = 3.41), similarly so in the vertical run (invalid M = 5.54%, SD = 3.56; valid: M = 3.21%, SD = 2.27). The BF_ANOVA for reaction times yielded strong evidence only for the main effect of cueing-condition with a BF_10_ of 17275.39 against the baseline model. This model was also superior to the other possible combinations of the 2×2 design (evidence in favor of the cueing only model against: direction only BF_10_ = 87028.43; both main effects BF_10_ = 5.05; main effects plus interaction BF_10_ = 15.7). The analyses of the error rates yielded similar results. The model including only a cueing main effect had the highest BF_10_ against the intercept model (BF_10_ = 16.8), and also stood out against all other possible combinations of factors (evidence in favor of cueing only against: direction only BF_10_ = 64.8; both main effects BF_10_ = 4.13; against main effects plus interaction BF_10_ = 8.32). In sum, these analyses show that the main manipulation of the experiment—the reorientation of attention in invalid trials—induced the expected reaction time costs and increased difficulty, as seen in the error rates. Moreover, they provided positive to strong evidence that neither the overall level of reaction times nor the reorienting costs after invalid cueing differed between the horizontal and vertical runs.

**Figure 3.**
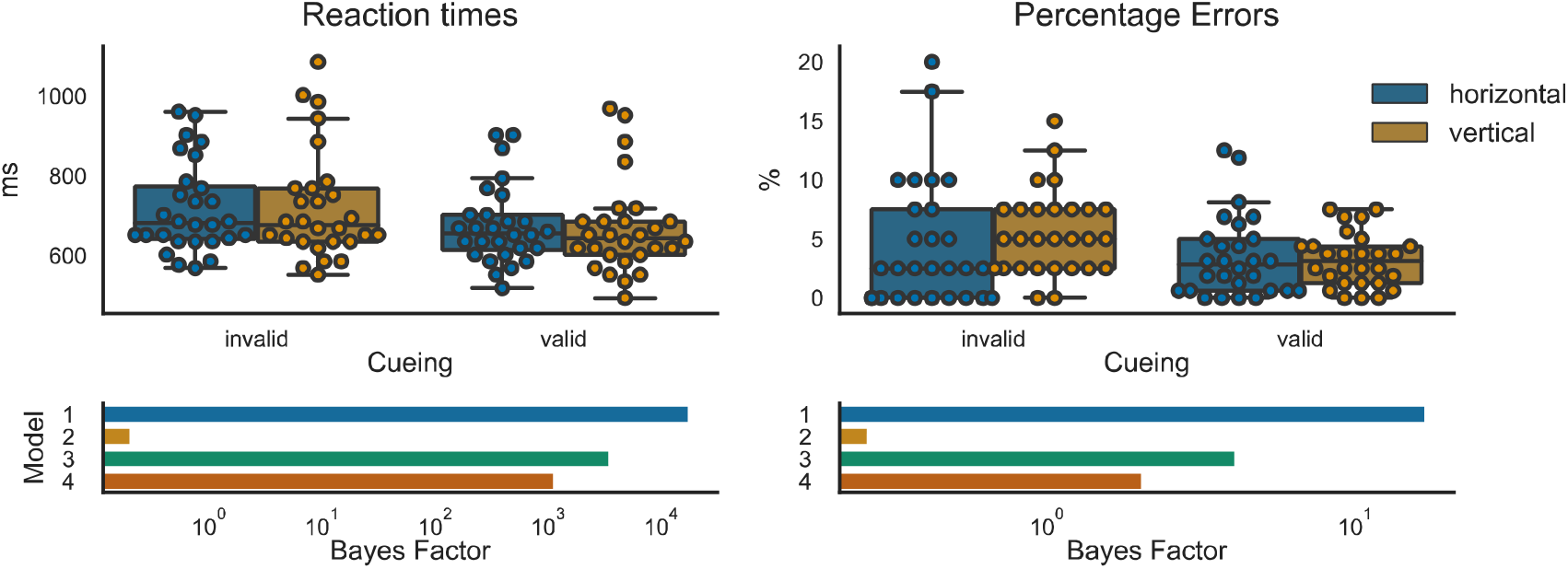
Results for the behavioral data. In the upper part of the figure, boxplots show the distribution of the data (median, IQR, and 90% percentile). Swarm plots were used to indicate individual data points in the sample. The bar graphs in the lower part indicate the Bayes factor (in logarithmic scale) against an intercept model. *Model1 - Cueing only; Model 2 - Direction only; Model 3 – Cueing + Direction; Model 4 – Cueing + Direction + Cueing * Direction*.

### 3.2 GLM

Figure 4 depicts the main effect of cueing (invalid > valid cueing, contrast (1)) for vertical and horizontal runs combined. The automatic calculation of the cluster-forming threshold at p < 0.001 (cluster corrected FWE p < 0.05) yielded a cluster forming threshold of k ≥ 58 voxels. Cluster size in cubic millimeter, global maxima, up to four local maxima, and their respective t-statistics are provided in Table 1. Reorienting across both runs activated areas of the dorsal and ventral frontoparietal attention networks. The largest cluster stretched along the parietal cortex, with the local maxima located in bilateral IPS and bilateral precuneus. The next cluster included the right FEF and extended into the right insular cortex, as well as into the medial and inferior frontal gyrus. In the right hemisphere, we found a single cluster in the TPJ. Similar activation patterns were observed in the left hemisphere, with separate clusters in the insular cortex, FEF, IFG, and TPJ.

**Figure 4.**
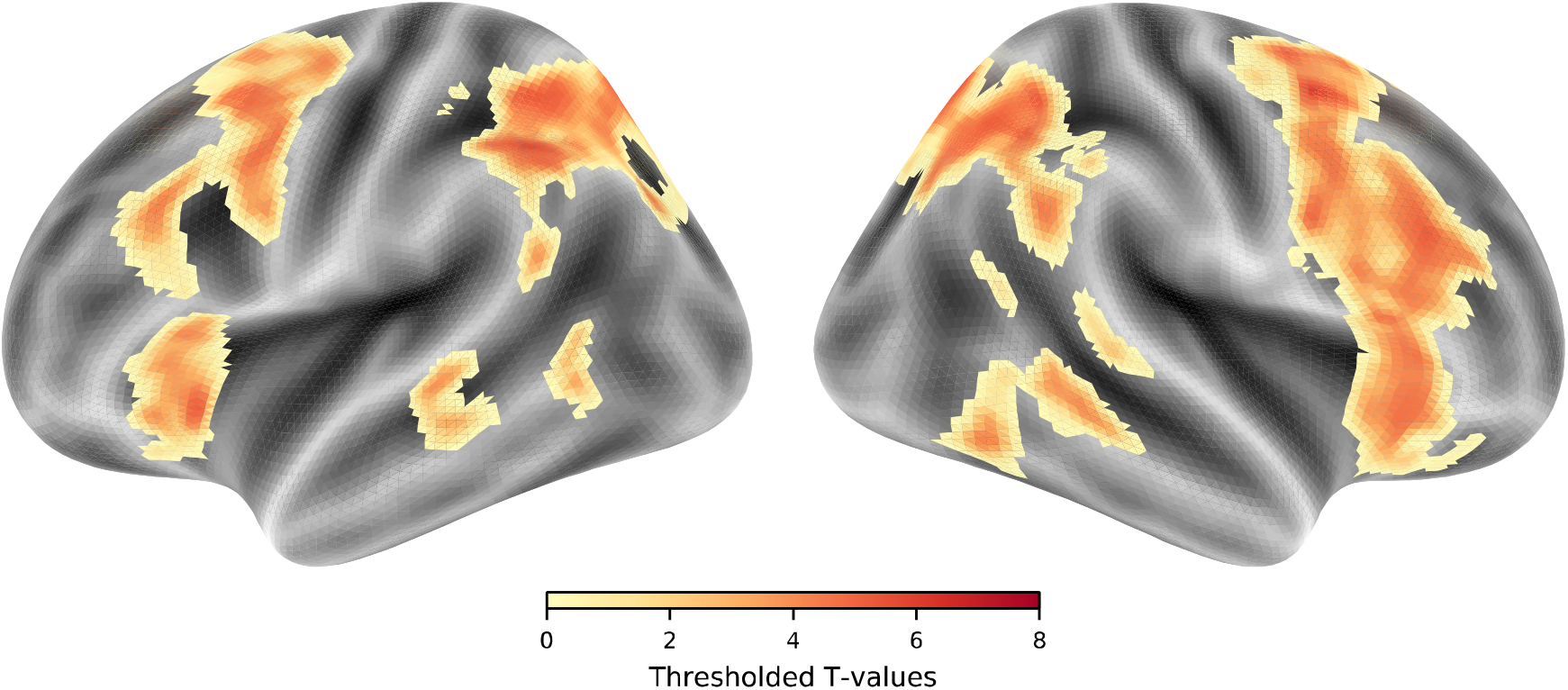
Statistical map for the reorienting (invalid > valid) across horizontal and vertical runs. The thresholded map was projected onto the freesurfer inflated surface templates (fsavg5) using nilearn.

The run-specific activation maps of reorienting-related activity are depicted in Figure 5. In the vertical run, clusters surviving the statistical threshold (k ≥ 57 voxels) were found in bilateral IPS and right TPJ. Additionally, significant activations were observed in the right inferior frontal and middle frontal areas, as well as in the insular cortex. The main effect of cueing in the horizontal run revealed clusters (k ≥ 47) in bilateral FEF and IPS.

**Figure 5.**
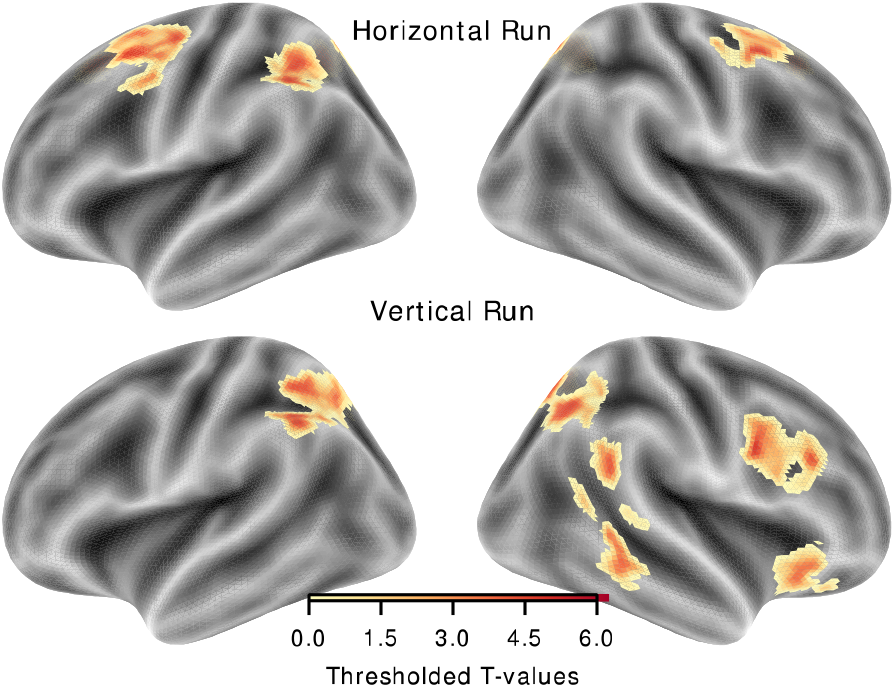
Statistical maps of the reorienting (invalid > valid) in each run. The thresholded maps for the two runs were projected onto the freesurfer inflated surface template (fsavg5 using nilearn.

Tests for main effects of direction (k ≥ 57) and the interaction of direction and cueing (k ≥ 47) did not yield any significant voxels surviving the cluster-based FWE correction.

**Table 2:** Cluster coordinates for reorienting-related activity in horizontal and vertical runs. Only the coordinates of the global maxima in the main clusters are reported. *FEF – frontal eye fields, SMA – supplementary motor area, TPJ – temporoparietal junction*.

| Global<br>Maximum | Side | MNI Coordinates |  |  | Peak<br>Statistic | Cluster<br>Size |
| --- | --- | --- | --- | --- | --- | --- |
|  |  | X | Y | Z | (T) | (mm) |
| Invalid > Valid Horizontal |  |  |  |  |  |  |
| Angular gyrus | R | 10 | -60 | 51 | 5.23 | 2578 |
| Parietal cortex | L | -43 | -51 | 47 | 5.12 | 4737 |
| FEF | L | -24 | 9 | 57 | 4.94 | 6413 |
| FEF | R | 29 | 6 | 51 | 4.89 | 2835 |
| SMA | L | -5 | 21 | 51 | 4.55 | 2964 |
| Precuneus | L | -12 | -60 | 51 | 4.53 | 2062 |
| Invalid > Valid Vertical |  |  |  |  |  |  |
| Precentral | R | 45 | 6 | 31 | 5.7 | 4382 |
| TPJ | R | 60 | -51 | 21 | 5.36 | 1869 |
| Insula | R | 29 | 27 | -12 | 5.03 | 2352 |
| Temporal | R | 63 | -35 | -9 | 4.89 | 2159 |
| Parietal cortex | L | -49 | -51 | 41 | 4.65 | 8153 |
| Parietal cortex | R | 20 | -73 | 54 | 4.56 | 9700 |
| Precuneus | L | -8 | -73 | 47 | 4.22 | 1353 |
| Lingual gyrus | L | -2 | -76 | 54 | 3.48 | 32 |

Our analysis of the attentional modulation in valid trials in relation to the visual fields (Figure 6) revealed activations (k ≥ 52) in left dorsal and ventral higher-order visual areas (including V4 and V5) for the contrast of right versus left valid targets. The reverse contrast (left versus right valid targets), yielded a cluster (k ≥ 53) of significant activation in ventral parts of right higher-order visual areas. Contrasting trials with lower visual field targets versus upper visual field valid targets resulted in a significant cluster (k ≥ 46) in right and dorsal parts of higher-order visual areas. The reverse contrast did not reveal any significant results.

**Figure 6.**
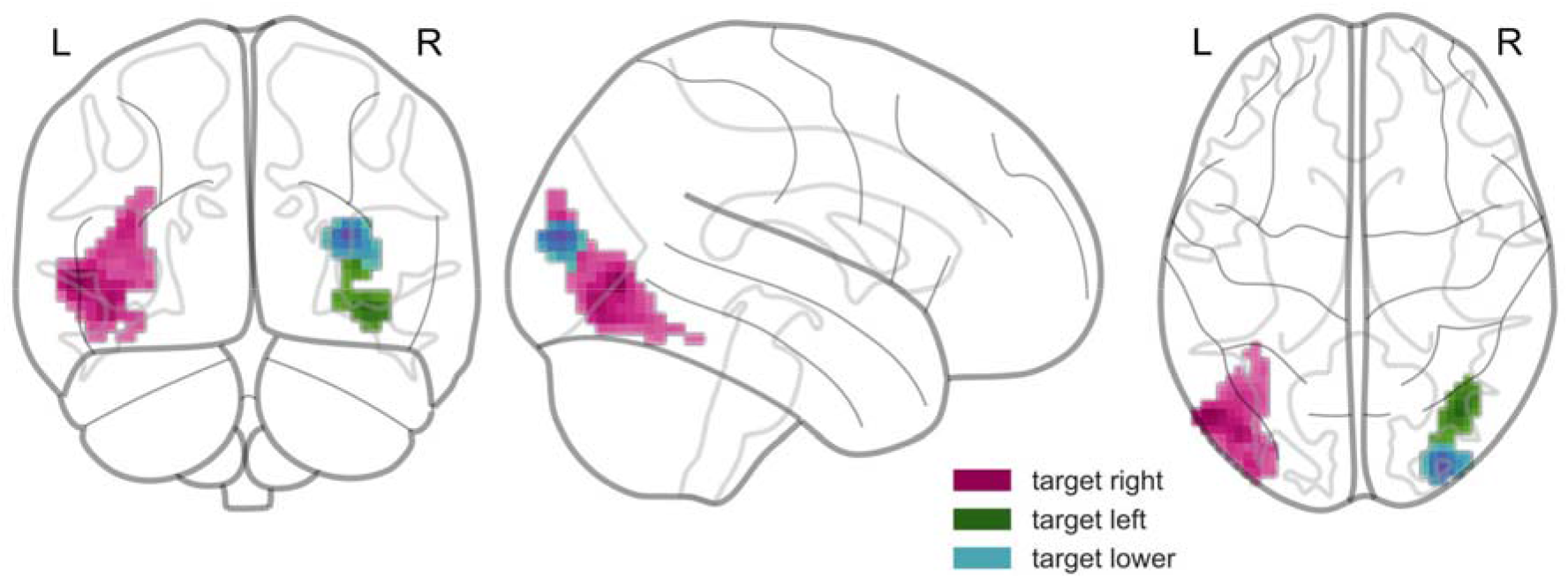
Attentional modulation by the direction of attention in valid trials (i.e., targets appearing in the left, right, upper, and lower visual fields).

The statistical t-maps of the GLM analysis can be found on Neurovault in a thresholded and un-thresholded form (https://neurovault.org/collections/NKRRQBJU/).

### 3.3 VOI Analyses

As a potentially more sensitive approach, we extracted the regression (beta) weights of the main GLM analysis in six regions that showed significant reorienting related activity (see Table 1). The BF_ANOVA (following the same rationale as in the behavioral analysis) yielded the highest evidence for the model including only the main effect of cueing in all six regions (BF_10_ for left IPS = 33.13; right IPS BF_10_ = 45.93; left FEF BF_10_ = 10.91; right FEF BF_10_ = 31.44; left TPJ BF_10_ = 5.92; right TPJ BF_10_ = 9.04). Comparing the cueing-only effect against the main effect of direction, both main effects, and main effects plus interaction (see Figure 7), showed that there was only positive evidence in favor of the cueing main effect (BF_10_ > 3) in most of the VOIs. In the right IPS VOI, however, there was only anecdotal evidence (BF_10_ = 1.07) favoring the cueing-only model against the model including both main effects, meaning that we cannot convincingly exclude an additional effect of direction for this region.

**Table 3:** Summary statistics VOI-based BF_ANOVA on the regression (beta) weights. Each row displays the mean (M) and standard deviation (SD) for each of the six VOIs. The BFs for the comparison of model 1 versus the three other models are shown. These BF_10_s indicate how much more likely model 1 is, compared to the other models. *Model1 - Cueing only; Model 2 - Direction only; Model 3 – Cueing + Direction; Model 4 – Cueing + Direction + Cueing * Direction*.

| VOI | Invalid |  | Valid |  | BF_ANOVA |  |  |
| --- | --- | --- | --- | --- | --- | --- | --- |
|  | Horizontal | Vertical | Horizontal | Vertical | Model 1 versus |  |  |
| | $M \pm SD$ | $M \pm SD$ | $M \pm SD$ | $M \pm SD$ | Model 2 | Model 3 | Model 4 |
| **IPS L** | 4.97 ± 3.60 | 5.19 ± 2.64 | 4.24 ± 2.93 | 4.22 ± 2.44 | 150.60 | 4.54 | 15.73 |
| **IPS R** | 3.27 ± 2.13 | 3.80 ± 1.96 | 2.69 ± 1.79 | 2.91 ± 1.63 | 61.47 | 1.07 | 3.24 |
| **FEF L** | 3.64 ± 2.10 | 3.53 ± 2.11 | 2.80 ± 1.77 | 2.90 ± 2.02 | 48.62 | 4.98 | 15.78 |
| **FEF R** | 3.07 ± 1.71 | 2.92 ± 1.67 | 2.47 ± 1.41 | 2.43 ± 1.51 | 135.10 | 4.42 | 15.35 |
| **TPJ L** | 1.60 ± 1.61 | 1.56 ± 1.66 | 1.02 ± 1.49 | 0.85 ± 1.54 | 26.72 | 4.51 | 15.31 |
| **TPJ R** | 2.10 ± 2.33 | 1.93 ± 1.86 | 1.35 ± 1.95 | 1.31 ± 1.74 | 39.99 | 4.36 | 16.10 |

**Figure 7.**
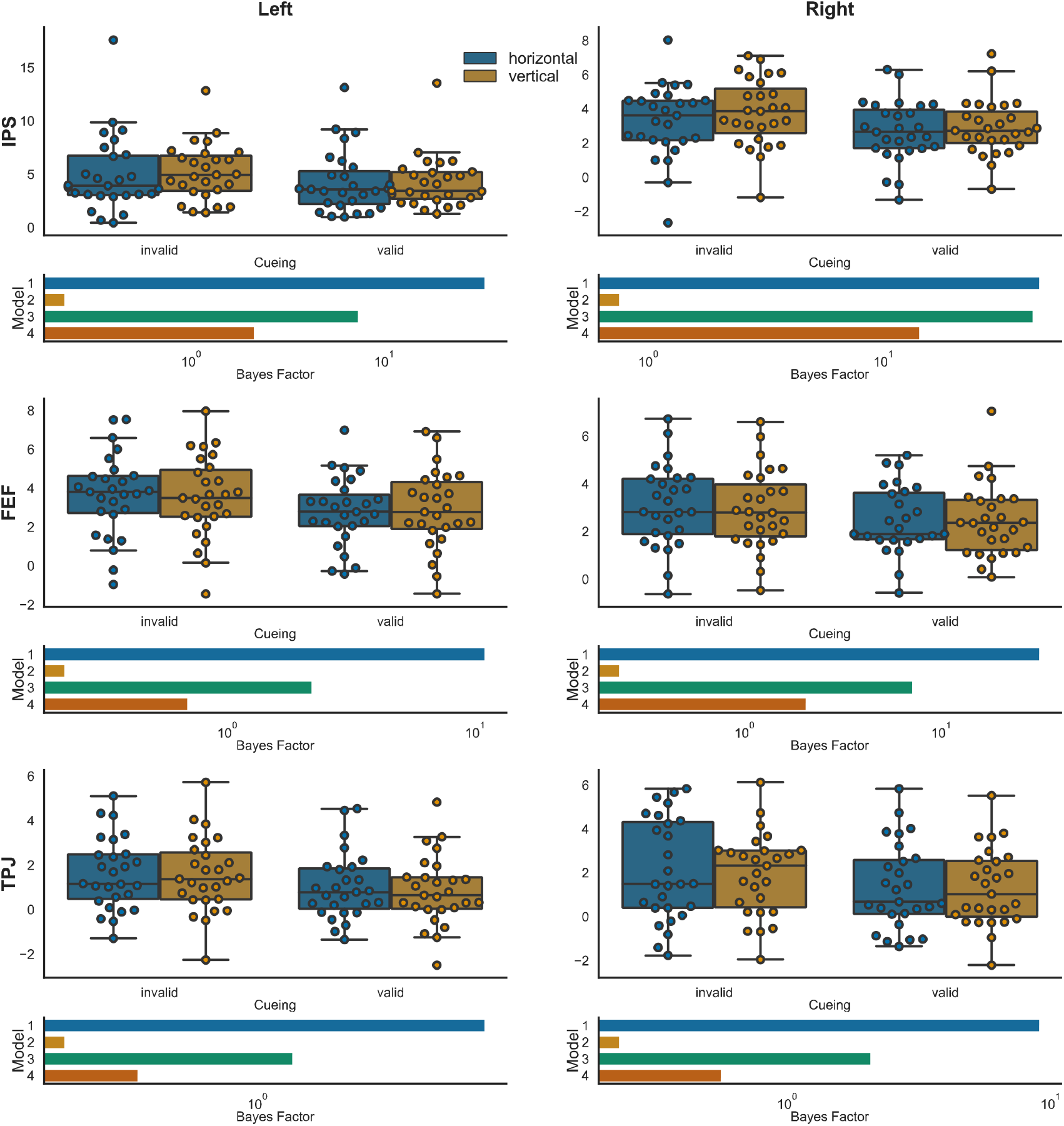
Results of the VOI based analysis. For each of the six VOIs, boxplots show the distribution of the data (median, IQR, and 90% percentile). Swarm plots were used to show individual data points in the sample. The bar graphs below the box plots indicate the Bayes factor (in logarithmic scale) against an intercept model. VOIs are displayed separately for left- and right-hemispheric regions and in the order IPS, FEF, TPJ. *Model1 - Cueing only; Model 2 - Direction only; Model 3 – Cueing + Direction; Model 4 – Cueing + Direction + Cueing * Direction.*

Using logistic regression, we tested whether the average beta weights of the eight regressors (valid and invalid trials for all target locations) could predict the cueing-condition (valid/invalid trails) in the respective other run. The prediction was significant for each run with an accuracy of 62.2% (P = 0.021) for the horizontal and with an accuracy of 62.3% (P = 0.021) for the vertical run. More importantly, the model trained on the horizontal run generalized to the vertical run with an accuracy of 59.3% (P = 0.016), and the model trained on the vertical run generalized to the horizontal run with an accuracy of 62.0% (P = 0.006). These results support the observation that the activation patterns in the six VOIs were highly similar, so that those predictive models generalized well across the two runs.

### 3.4 DCM Analyses

The DCM analysis was carried out using data from 26 of the remaining 27 participants, as for one participant, the coordinates for the left TPJ VOI could not be established. To select the DCM with the highest evidence of generating the network activity in our data, we applied a hierarchical familywise model selection. The family with modulations in both hemispheres was slightly superior (exceedance probability (eP) = 0.54) when compared to the other two families (left lateralization eP = 0.46, right lateralization eP = 0.00). This family was then further subdivided into three families consisting of models describing the direction of the interhemispheric IPS connections. The model family with a modulation from right IPS to left IPS had the highest evidence with an eP of 0.76 (eP IPS right to IPS left = 0.00; eP bidirectional modulation = 0.24). Finally, the models in this winning family were subjected to BMA. The participant-specific DCM models with averaged parameter estimates had a good to moderate fit to the data, with a mean coefficient of determination (R^2^) of 33.74 (SD = 10.39, range 16.65 to 63.33).

The parameters for invalid horizontal (B^1^) and invalid vertical (B^3^) trials were compared using Bayesian paired t-tests. Most modulations provided positive evidence for an absence of differences between both runs (Table 4). For the connections from left TPJ to left FEF, left FEF to left TPJ, and right FEF to right TPJ, there was only anecdotal evidence against a difference in parameters (M = 0.12, SD = 0.49, BF_10_ = 0.44; M = 0.13, SD = 0.47, BF_10_ = 0.48; M = −0.3, SD = 0.84, BF_10_ = 0.90).

**Table 4:** BMA parameter inference. Paired BF t-tests were employed to compare the differences in connection strength between invalid horizontal and invalid vertical trials based on the parameters resulting from the BMA. Mean (M) and standard deviations (SD) show the group paired statistics (i.e., horizontal – vertical parameters for each subject combination) and the BF_10_, indicating evidence for a difference between the two runs. Connections are ordered by left, right, and inter-hemispheric connections.

|  | Left Hemisphere |  |  |  | Right Hemisphere |  |  |  | Inter-hemispheric |
| --- | --- | --- | --- | --- | --- | --- | --- | --- | --- |
|  | TPJ to IPS | TPJ to FEF | IPS to TPJ | FEF to TPJ | TPJ to IPS | TPJ to FEF | IPS to TPJ | FEF to TPJ | IPS-R to IPS-L |
| **M ±** | 0.11 | 0.12 | 0.00 | 0.13 | -0.14 | 0.04 | 0.10 | -0.30 | -0.11 |
| **SD** | 0.96 | 0.49 | 0.43 | 0.47 | 0.94 | 0.45 | 0.61 | 0.84 | 0.98 |
| **$BF_{10}$** | 0.24 | 0.44 | 0.21 | 0.48 | 0.27 | 0.23 | 0.28 | 0.90 | 0.24 |

Further interrogating the DCMs for each participant revealed that DCMs based on the BMA (RMSE: M = 0.35, SD = 0.08) performed in general better than the random models (RMSE: M = 0.41, SD = 4.00, all P < 0.001, except for one with P = 0.006). Swapping the vertical and horizontal inputs (iH (u_1_) ↔ iV (u_3_), vH (u_2_) ↔ vV (u_4_)) led to slightly worse performance in each model (R^2^: M = 33.09, SD = 10.42), when compared to the original data. Still, the swapped model was superior to a random input model for most participants (RMSE: M = 0.37, SD = 0.09). Using a cutoff of P < 0.05 (i.e., 5% of random models had a lower RMSE than the swapped model), the model with swapped inputs performed better than the random input model in 18 out of 26 participants (69%).

## 4 Discussion

This fMRI study used two versions of a spatial cueing paradigm to compare the behavioral and neural mechanisms underlying attentional reorienting along the horizontal and vertical meridians. Regardless of cueing direction, our experimental procedures induced the well-established reaction time costs in responses following invalid cues when compared to valid cues (Hedge, Powell, & Sumner, 2017). Attentional reorienting behavior was comparable within participants for the vertical and the horizontal direction, suggesting that the costs of reorienting spatial attention are unaffected by directionality. Along the same lines, the analysis of the fMRI data using a GLM, a VOI-based approach, and DCM analyses revealed no evidence for direction-sensitive effects in the higher-level regions of the attentional networks.

To tackle the difficult task of quantifying the absence of an effect of direction or interaction of direction and cueing, we applied, wherever possible, Bayesian inference methods like Bayes factor ANOVAs and Bayesian t-tests to get an estimate of the likelihood of the presence (or absence) of the effects of interest. In the case of BF_ANOVAs, this provided us with the possibility to compare the main effect of cueing against other possible interactions and main effects. For the behavioral measures, this analysis revealed evidence in favor of a model including an effect of attentional reorienting only, without additional interactions. Similar results were observed in the fMRI analyses suggesting a similar neural mechanism of attentional reorienting in different spatial directions.

In addition to statistical analysis, we were also able to show that predictive models trained on BOLD data related to attentional reorienting along one meridian generalized well to the other. In other words, the effect of direction was not only statistically insignificant but also had no impact on the generalizability of statistical models - so that the cueing condition in one run could be successfully predicted by the model from the respective other run. This novel analysis approach, which does not rely on classical inferential statistics based on p-values, strongly suggests that the higher-order neural mechanisms underlying attentional reorienting are insensitive about different spatial directions.

Along the same lines, we also demonstrate that the network dynamics of a DCM between runs were so similar that they could be used to reproduce the BOLD activity patterns induced by attentional reorienting in the respective other spatial direction.

Our results replicate the findings of Macaluso and Patria (2007), who also did not find any significant differences between vertical and horizontal reorienting in a similar experimental set-up using classical inferential statistics. However, our study extends these findings in multiple ways since we considerably increased statistical power by including more than twice the number of participants in our study and employed the Bayesian and predictive approaches described above.

Still, other studies contrasting vertical and horizontal stimulus layouts have shown direction-sensitive effects for behavioral and neuroimaging data. For example, differential activity in superior parietal and frontal areas was found in fMRI studies using an attentional cueing paradigm (Mao et al., 2007), or vertical and horizontal saccades and anti-saccades (Lemos et al., 2016, 2017).

One reason for these discrepancies might be that horizontal and vertical asymmetries critically depend on the basic perceptual properties of the visual system. For example, it has been argued that horizontal and vertical asymmetries (Rizzolatti, Riggio, Dascola, & Umiltá, 1987) are particularly evident at high visual eccentricities (Abrams, Nizam, & Carrasco, 2012; Carrasco & Chang, 1995), where the different physiological properties of different parts of the retina become perceptually and behaviorally relevant (Carrasco, Talgar, & Cameron, 2001; Jóhannesson, Tagu, & Kristjánsson, 2018). In our current study, the stimuli were presented at relatively small eccentricities so that the stimulus configurations may have minimized the impact of early retinal asymmetries. At the same time, stimuli were located distant enough to allow for a specific attentional modulation in cortical visual areas, as indicated by selective functional modulations in response to valid target stimuli. Our experimental design controlled for early bottom-up influences and hence allowed determining cortical effects related to top-down control.

However, despite carefully controlling for bottom-up influences and using an attentional cueing task, Mao et al. (2007) reported a horizontal-vertical asymmetry in brain activity and behavior. A critical difference between this and our study concerns the informational value of the cues. In Mao’s study, the cues were always valid. Hence attentional reorienting could not be investigated. The cues in the present study were probabilistic (i.e., not always valid). This aspect is relevant for the level of uncertainty involved in attentional control since a higher level of uncertainty induces a preparedness for reallocation of visual attention. Eckstein, Shimozaki, and Abbey (2002) showed that perceptual properties of the target stimulus and its attentional enhancement do not modulate reorientation costs. Instead, expectations primarily drove them. This view is in line with other studies manipulating the percentage of cue validity in similar location-cueing paradigms and reporting effects on response times and brain activity in dorsal and ventral attentional networks. For instance, increased uncertainty during invalid trials increases activity in the ventral attention network (Vossel, Mathys, Stephan, & Friston, 2015; Vossel et al., 2012) and decreases activity in the dorsal network (Weissman & Prado, 2012).

Similarly, higher activity in the ventral network correlates with worse behavioral performance in valid trials (Wen, Yao, Liu, & Ding, 2012). It has been suggested that the ventral network, and particularly the right TPJ, seems to be more generally involved in tracking and updating of expectations. Moreover, there are stroke patients with lesions to the right TPJ who display impaired rule changing and belief updating behavior in non-spatial tasks (Danckert et al., 2012; Stöttinger et al., 2014; for a review on different TPJ involvements see Geng & Vossel, 2013). Hence, the processes critically related to attentional reorienting in the current study might not necessarily be location-specific but might represent higher-order functions such as the processing of expectancy violations.

While the previous studies focused on the right TPJ, we observed that invalid cueing heightens TPJ activation in both hemispheres. Similar bilateral involvement of the TPJ has been described previously (Beume et al., 2017; Macaluso & Patria, 2007; Silvetti et al., 2016). However, the exact functional role of left- and right-hemispheric areas in the ventral network might differ (Dugué, Merriam, Heeger, & Carrasco, 2018).

The spatial independence of attention networks observed in the present study seems to contradict clinical data: Patients with ventral parietal lesions to one hemisphere are not able to reorient attention to an invalidly or neutrally cued target in the visual field contralateral to the lesion (Posner, Walker, Friedrich, & Rafal, 1984). Since the ventral network is generally assumed to respond to invalid trials irrespective of the target hemifield, such behavior may reflect functional impairment of the dorsal system or dorsal-ventral interactions (Corbetta, Kincade, Lewis, Snyder, & Sapir, 2005). TMS studies provide strong evidence for spatially selective effects in the dorsal network. For example, a concurrent TMS-fMRI study, where participants attended to stimuli in the left or right visual field, showed that TMS over posterior parietal cortices could modulate activations in the contralateral extrastriate cortex (Blankenburg et al., 2010). Similarly, TMS over left or right FEF led to top-down modulation of ipsilateral extrastriate areas (Duecker, Formisano, & Sack, 2013; Silvanto, Lavie, & Walsh, 2006). Still, these effects may not be purely symmetric, as right IPS and FEF have been shown to modulate not only the contralateral, but also the ipsilateral visual areas in some studies (Sheremata & Silver, 2015; Silvanto et al., 2006). It should be noted that we did not find direction-specific activations in the dorsal network in the present study. However, bilateral stimulus displays have been found to mask direction-specific effects in the dorsal network (Molenberghs, Gillebert, Peeters, & Vandenberghe, 2008).

Unilateral lesions to the ventral system may, therefore, lead to dysfunction and imbalance in the reallocation of attention in the dorsal system, resulting in attentional deficits in the horizontal spatial dimension in patients with neglect (Corbetta & Shulman, 2011; Macaluso & Patria, 2007). The allocation and reorientation of attention along the vertical meridian, on the other hand, may be more robust to unilateral lesions, as a central stimulus display would be represented in both hemispheres. Following this line of thought, bilateral lesions should be necessary to cause altitudinal neglect, and this has indeed been observed in a few patients with bilateral lesions to temporal areas (Shelton et al., 1990) and parietal areas (Rapcsak, Cimino, & Heilman, 1988).

Further experiments will be necessary to investigate whether the dorsal and ventral attention network interact in the hypothesized way. Despite extensive work using fMRI, for example on the direction coding in IPS (Molenberghs et al., 2008; Vandenberghe et al., 2005), as well as attention-modulated receptive fields in the dorsal attention network (Sheremata & Silver, 2015), to date it remains to be determined whether directional coding can also be found in ventral parietal areas.

In conclusion, we observed that reorienting visuospatial attention along the horizontal and vertical meridians relies on very similar neural processes in frontoparietal areas of the dorsal and ventral attention network. The absence of direction-specific effects in the ventral attention network, together with the bilateral involvement of the TPJ, corroborates the notion that this network is involved in higher-order cognitive processes such as violations of prior expectations – rather than being dependent on stimulus properties, such as its spatial location (Geng & Vossel, 2013).

## Acknowledgements

SV was supported by funding from the Federal Ministry of Education and Research (BMBF, 01GQ1401). We also like to thank our colleagues at the INM-3 for their valuable feedback and suggestions throughout the development of the study and analyses.

## Data Availability Statement

The data that support the findings of this study are available on request from the corresponding author. The data are not publicly available due to privacy or ethical restrictions.

